# Streak-Aware Localization Microscopy Enables High-Throughput Brain Imaging Across Platforms

**DOI:** 10.64898/2026.03.04.709480

**Authors:** Xuyang Chang, Quanyu Zhou, Abigail Vigderman, Shuangyi Cheng, Yuanyang Guo, Lin Tang, Tian Jin, Chaim Glück, Junjin Yu, Baoyuan Zhang, Lukas Glandorf, Michael Reiss, Mohamad El Amki, Susanne Wegener, Xosé Luís Deán-Ben, Bruno Weber, Thomas Longden, Kailiang Xu, Liheng Bian, Zhenyue Chen, Daniel Razansky

**Affiliations:** Institute of Pharmacology and Toxicology, Faculty of Medicine, University of Zurich, Zurich, Switzerland; Institute for Biomedical Engineering, Department of Information Technology and Electrical Engineering, ETH Zurich, Zurich, Switzerland; State Key Laboratory of CNS/ATM & MIIT Key Laboratory of Complex-field Intelligent Sensing, Beijing Institute of Technology, Beijing, China; Program in Neuroscience, School of Medicine, University of Maryland Baltimore, Baltimore, MD, USA; Department of Pharmacology and Physiology, School of Medicine, University of Maryland Baltimore, Baltimore, MD, USA; College of Biomedical Engineering, Fudan University, Shanghai, China; Zurich Neuroscience Center, Zurich, Switzerland; Department of Neurology, University Hospital and University of Zurich, Zurich, Switzerland; Institute of Precision Optical Engineering, School of Physics Science and Engineering, Tongji University, Shanghai, China

## Abstract

Optical, ultrasound, and optoacoustic localization microscopy based on microparticle tracking has enabled surpassing the resolution limits imposed by ultrasound diffraction and optical diffusion in tissues. However, its reliance on high-speed (kilohertz) data acquisition systems for precise emitter localization and tracking substantially increases methodological complexity and data storage demands, limiting scalability and applicability beyond specialized benchtop platforms. Here, we present streak-aware localization microscopy (SALM) that employs localization- and tracking-free deep learning model to convert motion-blurred streaks originating from low frame rate recordings of flowing emitters into super-resolved structural and functional readouts. In optical implementations, SALM exploits streaks captured by low-speed cameras to recover capillary-level cerebrovascular maps across a variety of benchtop, miniaturized, and second near-infrared preclinical imaging platforms, achieving over 30-fold reduction in the reconstruction time compared to conventional localization pipelines. We further introduce three coded excitation strategies that embed finer time-varying vectorial flow signatures into individual streaks, enabling single-frame velocimetry and video-rate hemodynamic imaging. Extending SALM to ultrasound imaging enables high-fidelity vascular imaging with centimeter-scale penetration in rhesus macaque and rat brains while reducing plane-wave compounding frame rates by up to one order of magnitude. By overcoming long-standing trade-offs between spatiotemporal resolution and hardware complexity, SALM offers a flexible and scalable framework for next-generation super-resolution microscopy.

## Introduction

Large-scale cerebrovascular and neuronal networks typically exhibit fluctuations on the millisecond timescale ^[1–2]^. These rapid, spatially heterogeneous variations underlie normal cognition and constitute pathological fingerprints in multiple brain conditions such as stroke ^[3–4]^, Alzheimer’s disease ^[5]^, or brain injury ^[6–7]^. Capturing fleeting events demands imaging technologies that combine micron-scale spatial resolution with millisecond temporal precision, while spanning fields of view (FOV) large enough to link local microcirculatory changes to whole-brain states.

Contemporary neuroimaging techniques are typically implemented in either serial-scanning or parallel configuration. The former systems encompass two-/three-photon microscopy ^[8–9]^, optical coherence tomography ^[10]^, optoacoustic microscopy ^[11]^, as well as ultrasound imaging with sequential linear scanning ^[12]^, all of which employ serial beam scanning to effectively suppress spatial crosstalk and achieve lateral resolution approaching the corresponding optical or acoustic diffraction limit. However, the sequential acquisition imposes a rigid trade-off between frame rate and FOV, rendering real-time quantitative flow mapping across large-scale FOV impractical ^[13]^. In contrast, parallel imaging implementations, such as epifluorescence (widefield, WF) ^[14]^, laser speckle contrast imaging ^[15]^, plane-wave ultrasound imaging ^[16]^, and optoacoustic tomography ^[17–18]^, record signals from centimeter-scale FOV simultaneously at video rates. Nevertheless, these parallel strategies suffer resolution degradation due to optical scattering and the acoustic diffraction limit.

To visualize large-scale vascular networks while preserving capillary-level detail, super-resolution localization concepts that were originally introduced for superficial nanoscopy applications ^[19–20]^ have later been adapted to *in vivo* deep-tissue angiographic imaging with various modalities ^[21–26]^. These localization-based methods pinpoint and track individual emitters, unifying structural and functional (blood flow velocity and direction) readouts in a single paradigm. However, their capability comes with significant trade-offs in hardware demands and spatiotemporal resolution. Achieving frame-wise sparsity necessitates specialized detection system operating at kilohertz frame rates to accurately resolve the motion of densely distributed emitters with minimal inter-frame displacement ^[27]^. Moreover, reconstructing a single super-resolved image typically requires complex inter-frame analysis and the accumulation of tens of thousands of raw frames, which inflates data storage and computational demands. Additionally, using straight-line approximations to connect successive localizations can distort true motion trajectories. These limitations underscore an unmet need for large-scale, brain imaging capable of capturing both structure and function with high spatiotemporal resolution without imposing excessive technical complexity.

In this work, we introduce streak-aware localization microscopy (SALM), a generalizable approach for super-resolved structural and functional brain imaging across optical benchtop, miniaturized, second near-infrared (NIR-II), as well as ultrasound platforms. Unlike classical localization microscopy mechanism, which localizes and tracks individual emitters at kilohertz sampling rates to minimize motion blur, SALM marks a paradigm shift by leveraging the motion-blurred streak generated by each emitter under low sampling rates as a self-encoded spacetime barcode. In optical modalities, these streaks also introduce a new degree of freedom for temporally encoded illumination with designed trigger patterns (much like a mechanical “ticker-tape timer”), enabling time-stamping along the streaks and allowing single-frame velocimetry without the need for inter-frame analysis. To decode and interpret these streaks, we developed a localization- and tracking-free deep learning model (LTf-Net) and a spatiotemporal trajectory simulation engine. LTf-Net avoids the error accumulation of conventional localization-and-tracking pipelines at low frame rates, while the simulation engine provides realistic training pairs that enable effective *in vivo* generalization without ground-truth annotations. Overall, SALM eliminates the trade-off between resolution and hardware, making super-resolved brain imaging routine, portable, and scalable for both preclinical and clinical applications.

## Results

### Principle of SALM

Localization-based vascular imaging is typically achieved through localization and tracking of emitters flowing through the vasculature (Fig. 1a). Conventional approaches necessitate high-speed data acquisition to minimize inter-frame displacements (Fig. 1b), thereby imposing significant hardware constraints. SALM addresses this limitation by interpreting motion-blurred streaks as intrinsic trajectories via an end-to-end LTf-Net (Fig. 1c; Supplementary Video 1), which bypasses the error-prone localization and tracking inherent to traditional pipelines.

**Figure 1.**
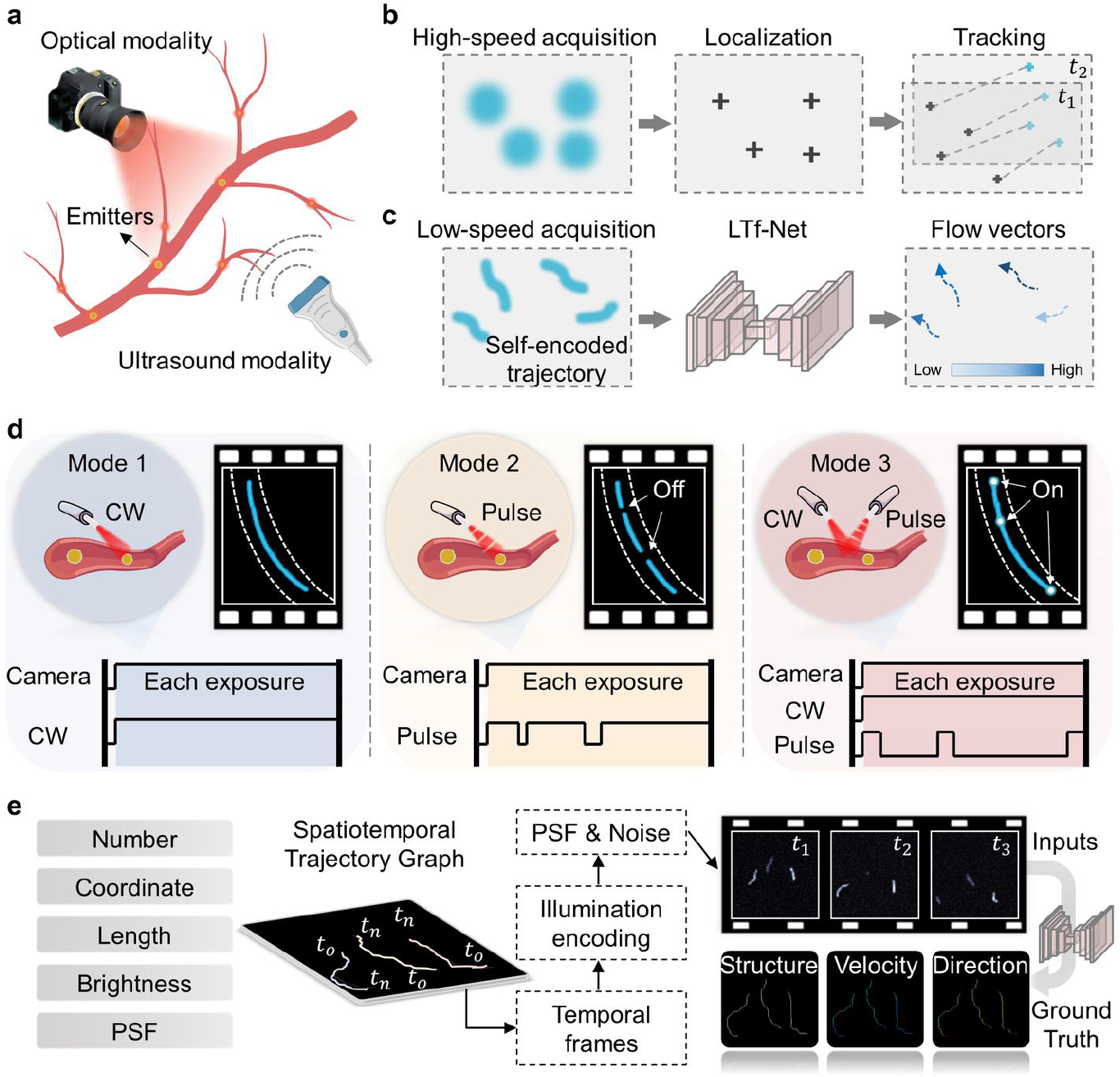
Principle of SALM. **a**, Scalability of SALM across optical and ultrasound modalities. **b**, Conventional localization microscopy mechanism requires high-speed acquisition to freeze motion for accurate tracking. **c**, SALM leverages motion-blurred streaks as signal carriers, eliminating the need for explicit localization and tracking via LTf-Net. **d**, Three illumination modes for optical-SALM embed vectorial flow information (velocity and direction) into each streak. **e**, Spatiotemporal graph-based simulation engine generates realistic paired datasets for network training.

The new mechanism makes SALM scalable across different imaging modalities. In the optical regime (Optical-SALM), motion streaks originate from continuous signal integration during exposure, granting high flexibility for excitation modulation. We further introduced three illumination schemes to embed temporal timestamps directly into streaks (Fig. 1d; Supplementary Note 1; Supplementary Video 2): Mode 1 employs continuous-wave (CW) illumination to produce uninterrupted streaks ideal for structural mapping, though functional recovery requires inter-frame analysis. Mode 2 utilizes pulsed excitation with unequal dark intervals acting as temporal fiducials, allowing simultaneous extraction of flow velocity and direction. Mode 3 superimposes pulses onto a continuous background to generate bright fiducial markers; this design preserves spatial continuity while uniquely enabling single-frame velocimetry for high-temporal-resolution functional imaging.

Unlike continuous excitation and detection used in Optical-SALM, ultrasound imaging typically operates in a pulse-echo mode, where short ultrasound pulses are transmitted and received, making it inherently immune to motion streaks within a single acquisition. However, state-of-the-art ultrasound localization microscopy (ULM) employs coherent plane-wave compounding to enhance spatial resolution and signal-to-noise ratio (SNR) ^[22]^. Hence, by taking advantage of motion streaks of flowing microbubbles during temporal integration of multi-angle transmissions, ultrasound-SALM can provide deep, non-invasive imaging capabilities while significantly lowering the frame rates, hardware complexity, and cost.

To recover vascular maps from the streaks recorded by SALM systems, we developed LTf-Net, integrating a U-Net encoder for spatial feature extraction with a long short-term memory (LSTM) module (Supplementary Note 2) ^[28–30]^. LTf-Net operates in an end-to-end manner, eliminating explicit localization and tracking, which are highly error-prone in low-frame-rate acquisitions. LTf-Net accepts either single-frame or multi-frame input and outputs super-resolved maps of vascular structure, pixel-wise estimates of flow velocity and direction. The simulation engine synthesizes spatiotemporal trajectories under excitation schemes, incorporating physical parameters such as physiological velocity range, point spread function (PSF), and noise level. This process produces paired datasets of streak images and corresponding ground truth (GT) maps for vascular structure and flow metrics (Fig. 1e; Supplementary Note 3).

### Phantom validation of optical-SALM

We first validate optical-SALM via phantom experiments, aiming to quantify the accuracy under three illumination modes (Fig. 2a; Supplementary Note 4). Microtubing with 280 µm inner diameter was used as a vascular-mimicking phantom. Laminar flow is generated by injecting fluorescent beads into the microtubing, with flow velocity controlled by a syringe pump to provide the GT. Given that cerebrovascular velocities in anesthetized mice range from 0 to 20 mm/s, we preset the bead velocities to 1, 5, 10, and 20 mm/s.

**Figure 2.**
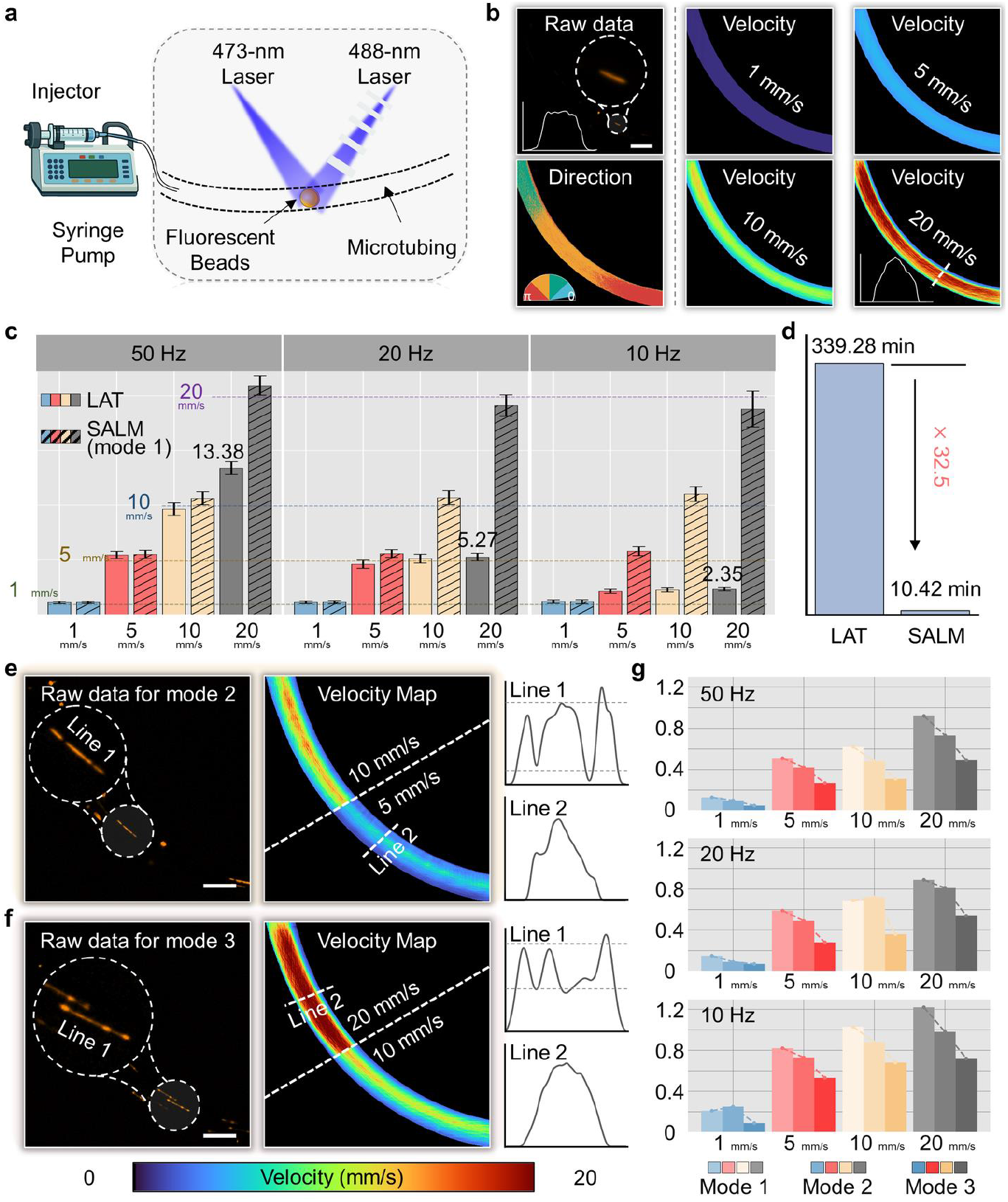
Phantom validation of optical-SALM. **a**, Experimental setup. **b**, SALM (mode 1) velocity estimation via motion streaks at 50 Hz. **c**, Quantitative velocity estimates from LAT and SALM (mode 1) across different frame rates and preset velocities. (dashed lines: preset values). **d**, Reconstruction times for LAT and SALM. **e-f**, Raw data and reconstructions for modes 2 and 3, showing temporal information embedding. **g**, Velocity estimation errors across all modes and conditions relative to preset GT. Scale bar: 500 µm.

Representative raw data and the corresponding intensity profile of a single bead from illumination mode 1 (top-left in Fig. 2b) demonstrate the generation of streaks using a video-rate camera (50 Hz). The reconstructed velocity maps closely match the preset GT and exhibit a near-parabolic profile (bottom-right in Fig. 2b), characteristic of laminar flow driven by a constant pressure gradient and balanced by viscous shear stress. The color-encoded direction map reveals gradual changes at the turns as expected (bottom-left in Fig. 2b). Quantitative evaluations of velocity accuracy are performed for both SALM (mode 1) and the conventional localization-and-tracking (LAT) method ^[31]^ (Supplementary Note 5) by computing the mean velocity within the microtubing. The maximum velocity resolvable by LAT declines rapidly with decreasing frame rate, averaging 13.38, 5.27, and 2.35 mm/s at 50, 20, and 10 Hz, respectively (Fig. 2c). In contrast, SALM maintained consistently high accuracy across all tested frame rates, demonstrating strong robustness to variations in camera frame rates. Additionally, the end-to-end LTf-Net substantially enhances computational efficiency by reducing the reconstruction time by 32.5-fold (Fig. 2d).

Streaked raw data were acquired under illumination mode 2 and mode 3 settings at 50 Hz, whereas the corresponding intensity profiles from individual beads reveal the embedding of temporal markers for vectorial flow, enabling finer velocity estimation within each streak (Figs. 2e-f). Absolute velocity errors of mode 1-3 across various frame rates and preset flow velocities demonstrate that mode 2 and mode 3 achieve higher accuracy, reducing estimation errors by 12.47% and 41.02%, respectively (Fig. 2g). These results underscore the effectiveness of illumination schemes in enhancing the accuracy of velocity estimation. Additional phantom experiments using looped microtubing further validate the performance of SALM in complex flow scenarios (Supplementary Note 6). In this setting, SALM successfully resolved velocity gradients induced by centrifugal effects, whereas LAT failed to capture these variations (Fig. S7d in Supplementary Note 6).

### Benchtop optical-SALM for super-resolved *in vivo* cerebrovascular imaging

Following phantom validations, we used the same benchtop platform but a different wavelength for *in vivo* cerebral vasculature imaging in mice post craniotomy (Fig. 3a). The paired 640 nm (pulsed mode) and 660 nm (CW mode) lasers were selected to form motion streaks of DiD-labeled red blood cells (RBCs) (Supplementary Note 7) ^[32]^. A camera records raw frames at 50 Hz.

**Figure 3.**
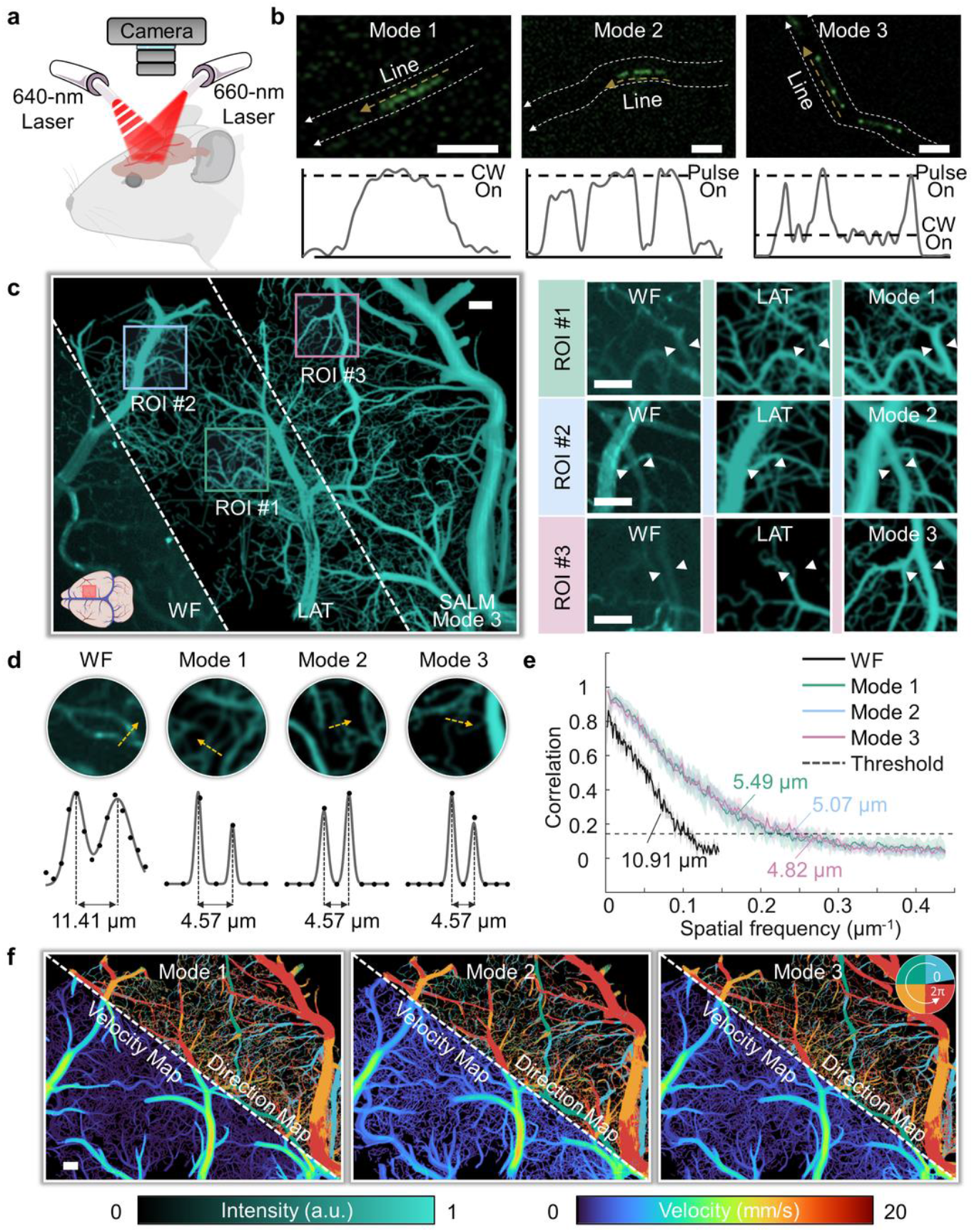
Benchtop optical-SALM for super-resolved *in vivo* cerebrovascular imaging. **a**, Experimental setup. **b**, Visualization of *in vivo* raw data acquired using three illumination modes. **c**, Comparison of cerebrovascular structural maps. **d**, Resolution assessment via double-Gaussian peak fitting. **e**, Quantitative resolution assessment using Fourier ring correlation. **f**, Super-resolved blood flow vector maps (velocity and direction) rendered from mode 1-3. Scale bar: 100 µm.

Raw images and their intensity profiles along individual stained RBCs rendered with the different illumination modes of SALM demonstrate reliable streak encoding despite intense photon scattering in living tissues (Fig. 3b). By integrating 200 s of data, we reconstructed three-fold super-resolved structural maps (Fig. 3c). The composite view compares WF image, LAT, and SALM (mode 3). WF resolved only the larger vessels due to the optical diffraction limit and photon scattering, whereas both LAT and SALM images exhibited enhanced spatial detail to resolve capillaries benefiting from super-resolved mechanism. When comparing with SALM, we observed some unresolved vessels in LAT images, as expected due to tracking failures in vessels with high flow velocities (indicated by white arrows in the regions of interest, ROIs). In contrast, SALM renders high-fidelity vascular structures that clearly delineate both micron-scale capillaries and larger arteriole/venules.

The achieved spatial resolution is further quantified by the peak-to-peak separation of adjacent capillaries (Fig. 3d; Supplementary Note 8), which shows a 2.49-fold improvement (4.57 µm) compared to WF imaging (11.41 µm). Fourier ring correlation (FRC) ^[33]^ analysis yields consistent results (Fig. 3e; Supplementary Note 8). The resolution limit of SALM is primarily determined by the SNR and camera pixel size, similar to conventional localization microscopy ^[34]^. The reconstructed velocity and direction maps maintain the same spatial resolution as the structural images (Fig. 3f; Supplementary Note 9). In addition, we evaluated imaging performance in a transcranial configuration, where SALM exhibited strong resilience to skull scattering and enabled clear visualization of high-flow arterial structures beyond LAT (Extended Data Fig. 2). FRC analysis yields a spatial resolution of 7.58 µm, representing a moderate reduction compared to the 4.82 µm resolution achieved with a cranial window. Structural results from SALM (mode 3) and LAT using different numbers of accumulated frames demonstrate the superior signal detection efficiency of SALM, enabling reliable results with shorter acquisition times (Extended Data Fig. 3; Supplementary Video 3).

To benchmark the *in vivo* performance of SALM against the absolute GT, we generated temporally downsampled datasets using LAT reconstructions from high-speed recordings as the GT (Extended Data Figs. 4–6). At the same frame rate, SALM achieves 8.87 dB improvement (48.95%) in peak signal-to-noise ratio (PSNR) and 0.166 improvement (25.15%) in structural similarity index measure (SSIM) compared to LAT, averaged across structural, velocity, and direction maps. Among the variants, illumination mode 2 and mode 3 demonstrate higher PSNR and SSIM than mode 1, with mode 3 showing the better performance under high-density emitter conditions.

**Figure 4.**
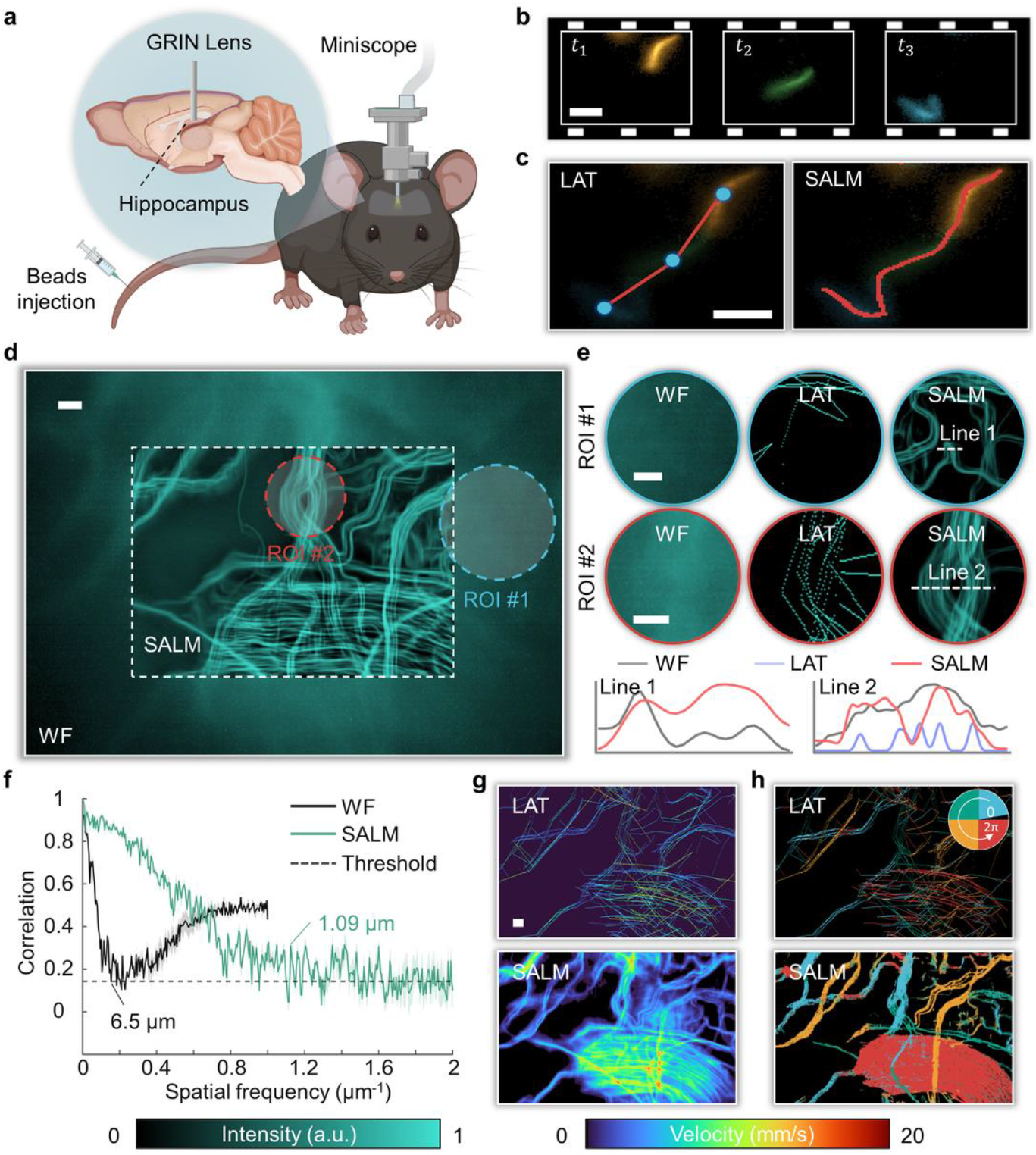
Miniaturized optical-SALM for hippocampal imaging in awake mice. **a**, Experimental setup via a UCLA Miniscope. **b**, Representative raw data across three consecutive frames. **c**, Mechanism comparison between LAT and SALM. **d**, Structural maps acquired with WF imaging versus SALM, demonstrating enhanced structural detail and improved SNR. **e**, ROIs and intensity profiles of different methods. **f**, Quantitative resolution using FRC. **g-h**, Comparison of velocity and direction maps reconstructed by LAT and SALM, respectively. Scale bar: 25 µm.

### Miniaturized optical-SALM for hippocampal imaging in awake mice

Miniaturized microscopes enable recording of brain activity in awake or freely moving animals while still preserving their natural behavior ^[35–36]^. However, the limited acquisition frame rate and relatively low SNR become dominant constraints in these platforms due to the strict requirement for sensor compactness, making quantitative blood flow imaging particularly challenging. To address this, we integrated SALM with the UCLA miniscope ^[37]^ and applied it for deep hippocampal imaging in awake mice through a chronically implanted GRIN lens (Fig. 4a; Supplementary Note 10).

Representative raw recordings of three consecutive frames exhibit motion-induced streaks (Fig. 4b) due to the limited frame rate of 30 Hz. Visualization of the resolved trajectories further illustrates the fundamental mechanistic differences between LAT and SALM (Fig. 4c). LAT localizes, tracks, and connects emitters across frames using straight-line approximations, which can distort vascular anatomy. In contrast, SALM directly interprets motion streaks to reconstruct true vascular architecture. The structural maps indicate that SALM offers higher resolution than WF imaging and greater morphological fidelity than LAT (Figs. 4d-e). FRC analysis (Fig. 4f) confirms a ~6-fold resolution improvement over WF (1.09 µm vs. 6.5 µm). Moreover, SALM achieves higher precision than LAT in functional readouts, including velocity and direction measurements (Figs. 4g-h). SALM thus unlocks super-resolved and high-SNR imaging on miniaturized platforms without additional hardware modifications, providing a promising platform for investigating complex hippocampal functions in awake mice.

### NIR-II optical-SALM for super-resolved, non-invasive cerebrovascular imaging

Although both benchtop and miniatured SALM platforms provide capillary-level resolution, these methods still require invasive surgeries for optimal imaging performance. Recent advances in NIR-II (1000-1700 nm) imaging have enabled deeper penetration due to reduced scattering and autofluorescence ^[38]^. However, most short-wave infrared (SWIR) sensors operate below 100 Hz, limiting accurate particle tracking for localization-based imaging. To achieve non-invasive super-resolved imaging, we developed a NIR-II SALM (Fig. 5a). We imaged the murine brain noninvasively at 18 Hz following intravenous injection of microdroplets encapsulating core/shell PbS/CdS quantum dots (Fig. 5b; Supplementary Note 11).

**Figure 5.**
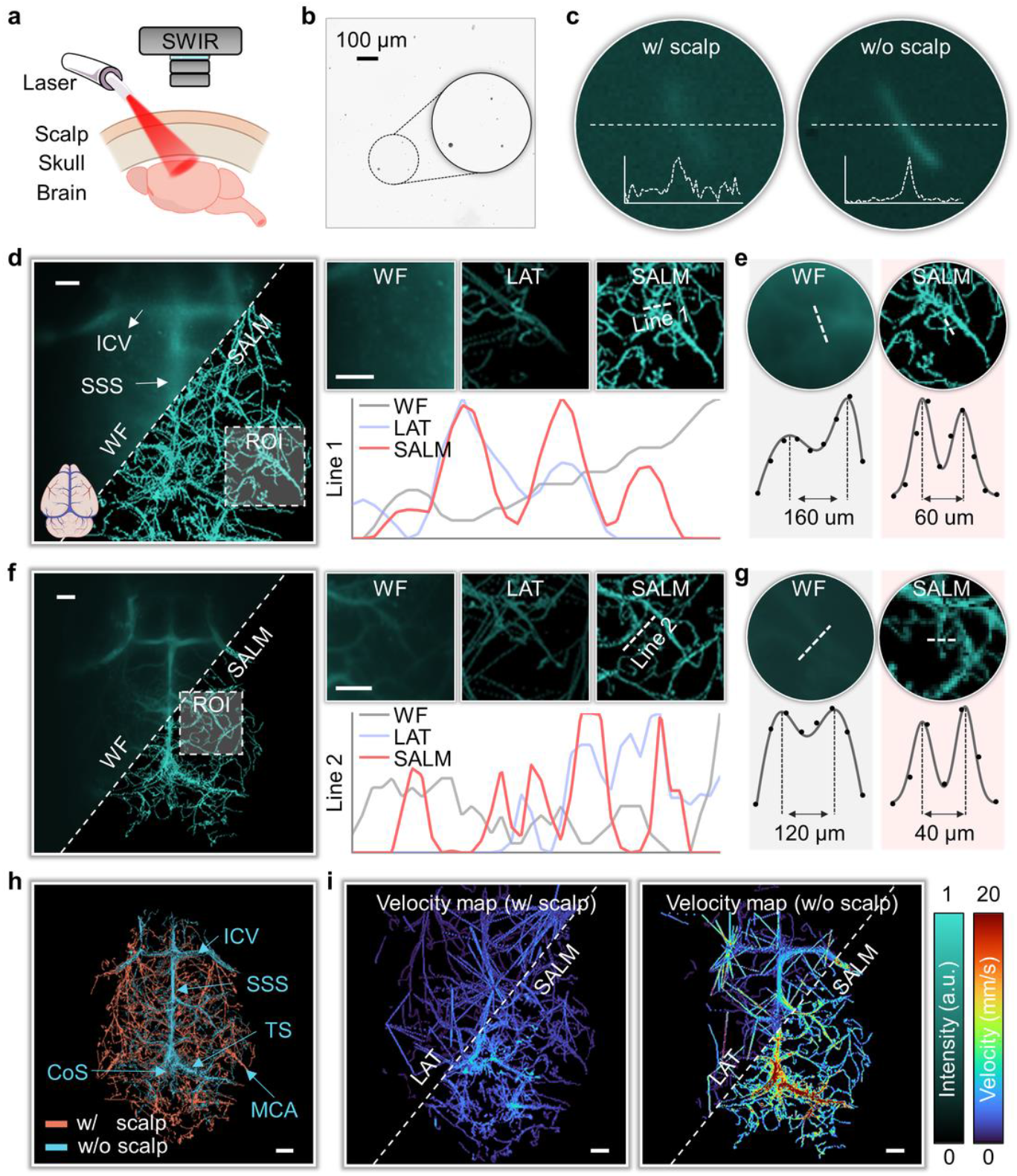
NIR-II optical-SALM for non-invasive imaging through intact scalp and skull. **a**, Experimental setup. **b**, Microdroplets inspection (~5-15 µm size). **c**, Raw data with vs. without scalp. **d-g**, Structural maps (with zoomed ROIs/line profiles) and resolution assessments acquired with (**d, e**) and without (**f, g**) intact scalp. **h**, Merged structural map. **i**, LAT and SALM velocity map comparison. ICV, inferior cerebral vein; SSS, superior sagittal sinus; MCA, middle cerebral artery; TS, transverse sinus; Cos, confluence of sinuses. Scale bar: 1 mm.

Representative raw streak images demonstrate the deep-tissue detection capability of NIR-II SALM (Fig. 5c). Scalp-induced scattering reduces SNR, challenging the signal detection compared to transcranial conditions. As a result, the WF image (Fig. 5d) resolves only major vessels, such as the superior sagittal sinus (SSS) and inferior cerebral vein (ICV). In contrast, SALM produced super-resolved structural maps that clearly delineated smaller arterioles and venules, as highlighted in the ROIs and intensity profiles (right, Fig. 5d). Resolution quantification via peak-to-peak separation confirmed more than a 2-fold improvement of SALM over WF imaging (Fig. 5e). The results after removing the scalp also confirm the superiority of SALM (Figs. 5f-g), and result in a fused image for spatial differentiation between scalp and cerebral vasculature (Fig. 5h). Velocity maps revealed the expected flow hierarchy, with fast velocities in major cortical vessels and slower flow in the superficial scalp vasculature (Fig. 5i).

### Single-frame optical-SALM for video-rate hemodynamic imaging

Resolving rapid changes in cerebral blood flow is essential for functional brain studies. However, forming a compounded localization image requires the temporal accumulation from thousands of raw frames, limiting the effective temporal resolution to 0.5-1 Hz ^[22]^. We demonstrate that SALM offers a promising approach to overcome the intrinsic trade-off between temporal and spatial resolution by enabling snapshot velocimetry without inter-frame analysis, and we further apply this capability to functional imaging in mice following hindpaw stimulation. The experiments were performed using the benchtop SALM (mode 3) operating at a 50 Hz frame rate (Fig. 6a). Six repeated stimulation cycles were applied, with detailed timelines shown in Fig. 6b.

**Figure 6.**
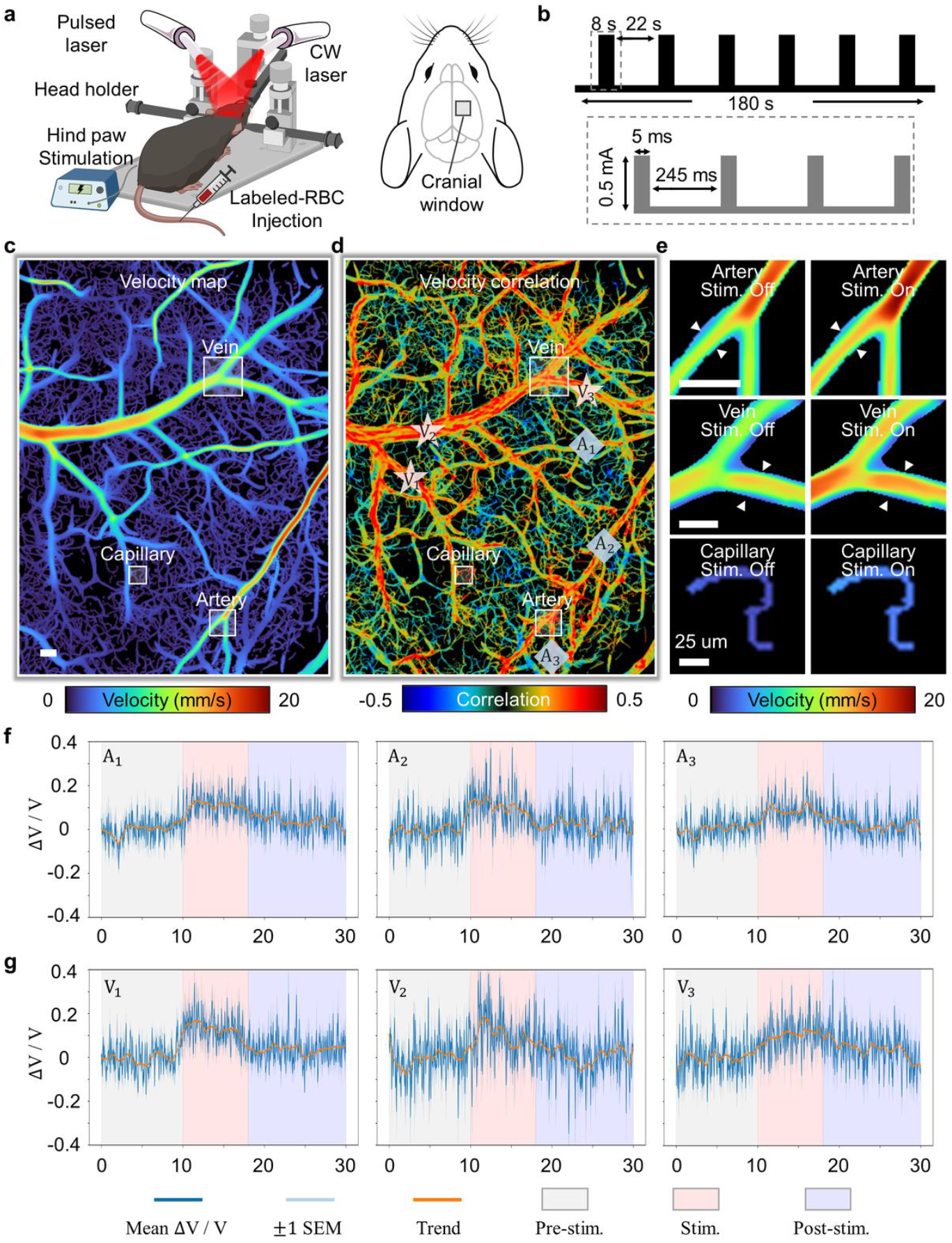
Single-frame optical-SALM for video-rate hemodynamic imaging. **a**, Experimental setup and the imaged region. **b**, Electrical stimulation module. **c**, Super-resolved velocity map obtained by accumulating all frames. **d**, Velocity correlation map identifying vessels with strong stimulus-evoked responses. **e**, Visualization of velocity changes in selected vessels (indicated in **c**) by averaging velocity maps from non-stimulated and stimulated periods. **f-g**, Video-rate (50Hz) response curves of the selected arteries and veins (indicated in **d**) from six stimulation cycles. Stim., stimulation; SEM, standard error of the mean. Scale bar: 100 µm.

The velocity map stacks were first reconstructed at a temporal resolution of 1 Hz, with the averaged map shown in Fig. 6c. The sensory-triggered hemodynamic activation pattern was characterized by calculating the correlation between the velocity profiles and stimulation pattern (Fig. 6d; Supplementary Note 12). Guided by the correlation map, we selected representative vessels with high correlation coefficients for further analysis. Velocity maps of selected arteries, veins, and capillaries (white rectangular box in Fig. 6c) were averaged across trials, revealing a clear stimulus-evoked increase during the stimulation period compared with the baseline (white arrows in Fig. 6e). We next performed single-frame inference on selected artery branches (rhombuses in Fig. 6d) and vein branches (stars in Fig. 6d) at the native acquisition rate of 50 Hz. The resulting high-temporal-resolution velocity response curves (Figs. 6f-g; Supplementary Note 12; Supplementary Video 4) reveal rapid increases in flow velocity in both arteries and veins following stimulus onset. These findings align with the expected vasodilation and enhanced blood flow associated with neurovascular coupling ^[39]^.

Quantitative validation of stimulus-evoked responses and amplitudes was conducted using temporally downsampled data (Extended Data Fig. 7). Raw recordings were acquired with a high-speed camera at 800 Hz and subsequently downsampled to 40 Hz. The velocity maps reconstructed by various SALM modes (Extended Data Fig. 7a) closely resembled the GT. Quantitative metrics (PSNR and SSIM) further confirmed the high fidelity of the reconstructed maps. The fitted trends of high-temporal-resolution response curves (40 Hz), extracted from selected arteries and veins, demonstrate that illumination mode 3 offers superior single-frame inference performance with higher accuracy in both temporal dynamics and response amplitude (Extended Data Figs. 7b-c).

### Ultrasound-SALM for deep, transcranial imaging of nonhuman primate

While ULM has enabled super-resolution microvascular imaging in deep tissue ^[21–22]^, its translational potential is hampered by skull-induced aberrations and highly demanding requirements for ultrafast frame rate acquisitions, both contributing to the system’s complexity, degraded SNR, as well as compromised microbubble (MB) localization and tracking performance when employing conventional acquisition and processing pipelines ^[40]^. Herein, we further extended SALM to low-frame-rate ULM and showcase its performance for brain imaging in a nonhuman primate (NHP, rhesus macaque) and rat models, the former offering critical translational insights due to its physiological and structural similarities to humans ^[41–42]^.

We performed ULM in an adult rhesus macaque using a phased-array transducer with diverging-wave acquisition in the sagittal brain plane (Fig. 7a). Unlike optical-SALM, ultrasound motion streaks arise from the temporal summation across multi-angle compounding (Fig. 7b). Performance was evaluated at effective frame rates of 400 Hz and 200 Hz, with 1000 Hz data serving as the GT. When comparing with the GT, the combined effects of lower SNR and temporal decorrelation of MBs at these reduced rates caused the conventional LAT processing pipeline to fail, producing artifactual microvascular branches (Fig. 7c, zoom-in views) and nonphysical velocity discontinuities (Figs. 7c-d). In contrast, SALM minimized the error accumulation, recovering realistic microvascular topologies with velocity profiles that closely matched the GT. Quantitative assessment using FRC confirmed that SALM achieved spatial resolutions of 161.8 µm (400 Hz) and 171.2 µm (200 Hz) (Fig. 7e), outperforming LAT (168.8 μm and 184.6 μm). Furthermore, velocity distribution histograms revealed that LAT significantly underestimated flow speeds, with deviations of 39.9% (400 Hz) and 52.9% (200 Hz) in peak velocities relative to the GT. In contrast, SALM effectively mitigated this underestimation, limiting the deviations to 21.7% and 39.9%, respectively (Figs. 7f-g).

**Figure 7.**
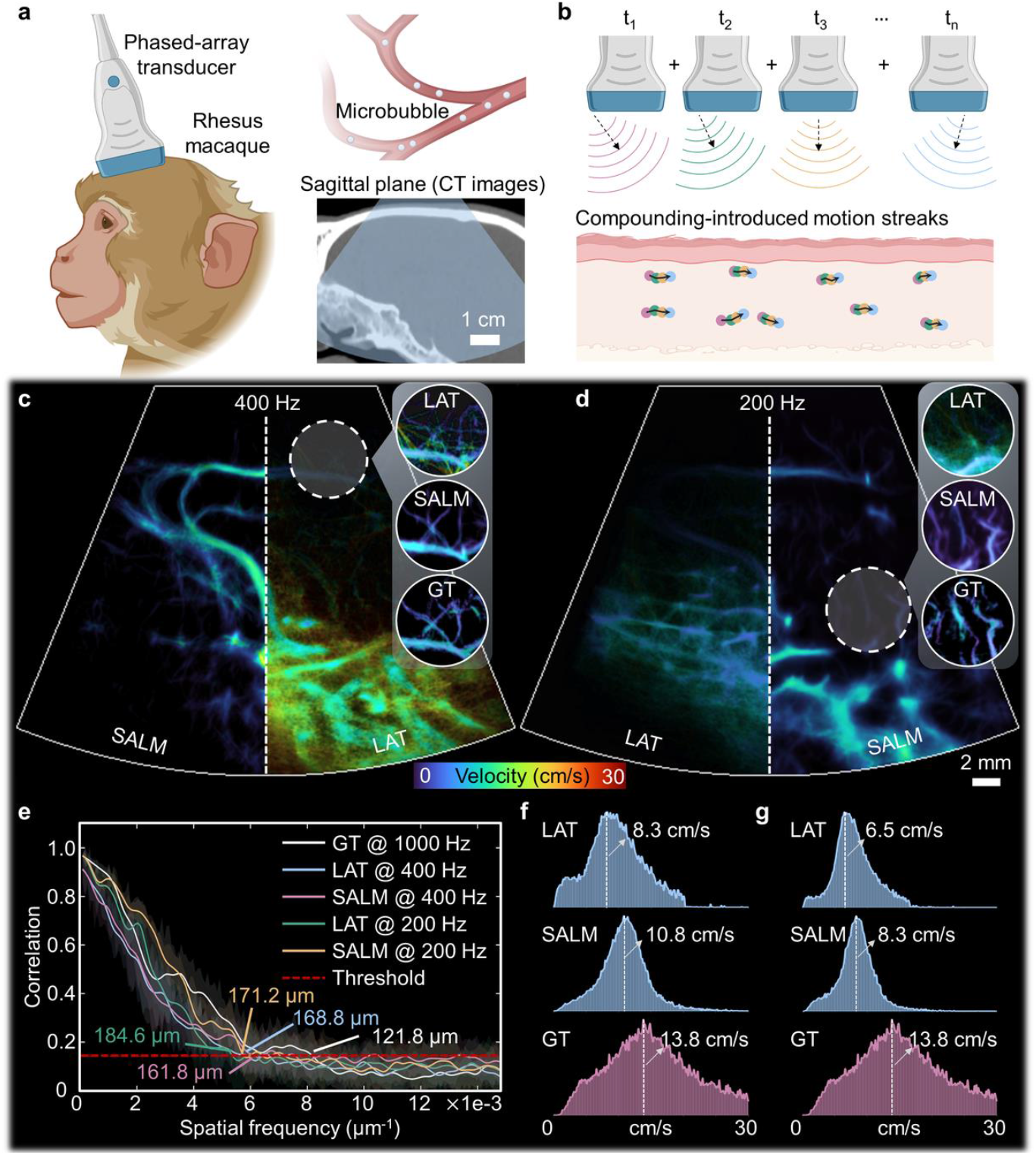
Ultrasound-SALM for deep, transcranial imaging of nonhuman primate. **a**, Schematic of the transcranial ULM setup in a Rhesus macaque. **b**, Principle of streak generation in ULM. Ultrasound streaks result from the coherent compounding of multiple wave emissions over time. **c**, Comparison of super-resolution velocity maps at an effective frame rate of 400 Hz between LAT, SALM, and the 1000 Hz GT. **d**, Velocity map comparison at a reduced frame rate of 200 Hz. **e**, Quantitative resolution assessment using FRC across different methods and frame rates. **f-g**, Histograms of flow velocity distribution at 400 Hz (f) and 200 Hz (g).

To further investigate the lowest possible frame-rates where the ultrasound-SALM method remains effective, we conducted experiments in the rat brain following craniotomy, which provides ideal high-SNR conditions. Axial velocity-coded maps with diverging colormap, rendered at effective frame rates of 75-150 Hz, showed that SALM preserved finer vascular morphology and more consistent flow patterns (Extended Data Figs. 8a-b). The reduced compounding frame rate led to a decreased SNR, thereby reducing MB detectability. Consequently, LAT reconstructions became sparse under the same accumulation time, whereas SALM maintained robust vessel connectivity (Extended Data Figs. 8c-d). Velocity histograms further confirmed the high accuracy of SALM in retrieving the hemodynamic information (Extended Data Figs. 8e-f).

## Discussion

Recent advances in neuroscience underscore a growing need for imaging tools that offer high spatiotemporal resolution and can be adapted across diverse platforms. However, existing approaches such as serial-scanning, widefield, and localization-based methods often involve hard trade-offs between various performance metrics and typically rely on specialized and expensive hardware, limiting their portability and broader applicability. In this work, we present SALM, which preserves the advantages of super-resolved emitter-localization methods while dispensing with high-end instrumentation. This allows for seamless deployment across various platforms, including optical benchtop, miniaturized, NIR-II, as well as ultrasound systems. Optical-SALM is further enhanced through an illumination encoding strategy that embeds temporal information directly into streaks within single frames. This bypasses the need for complex inter-frame analysis in conventional localization microscopy, enabling single-frame velocimetry with substantially improved temporal resolution for functional imaging. We further developed LTfNet that extracts spatial and temporal features directly from raw streak data, avoiding tracking-error accumulation at low frame rates and reducing computation time by more than 30-fold. Training is facilitated by a realistic simulation engine that generates paired datasets, ensuring robust generalization to *in vivo* conditions.

Despite these advances, SALM has certain limitations. First, although optical-SALM (mode 3) supports single-frame velocimetry, it requires a specific frame rate range to generate streaks long enough for temporal encoding, resulting in suboptimal sensitivity to slow flow. Future extensions could involve PSF-based encoding strategies that directly reshape the spread morphology of low-speed beads in the captured images to distinguish both velocity and direction ^[43]^. Second, the SNR affects the super-resolved reconstruction and the accuracy of velocity and direction mapping in SALM, as it does in conventional localization microscopy ^[34]^. As a result, SALM performance tends to degrade on miniaturized or other low-SNR platforms. Emerging deep optics approaches, which jointly optimize imaging hardware and reconstruction algorithms, offer a promising solution ^[44]^. By co-designing the imaging components and LTf-Net, the robustness of SALM under low-SNR conditions can be improved. Third, the present simulation framework is rooted in graphics-oriented rendering paradigms and therefore departs from authentic hemodynamic behavior. Integrating anatomically faithful vascular morphologies with physiologically realistic flow boundary conditions and rheological parameters would more accurately emulate *in vivo* hemodynamics ^[45]^, yield higher-fidelity training data, and ultimately enhance the network’s ability to transfer to experimental measurements.

The versatility of SALM has been demonstrated across several modalities and platforms. Optical-benchtop implementations provide capillary-scale, super-resolved neurovascular imaging *in vivo* and snapshot functional recordings at video rate. Miniaturized systems allow hippocampal imaging in awake mice, while NIR-II systems achieve fully noninvasive cortical imaging. Extended ultrasound-SALM addresses the challenges of low-frame-rate clinical systems, achieving centimeter-scale penetration depth in NHP. Looking forward, the developed conceptual framework is broadly transferable. In optical nanoscopy, SALM could exploit intrinsic sample drift to map sub-cellular dynamics and enable long-term live-cell imaging ^[46]^. Optical-SALM could also be extended to 3D acquisition using cylindrical lenses ^[47]^ or other PSF engineering methods ^[48–49]^, enabling volumetric flow field reconstruction with minimal loss in spatial or temporal resolution. From an application perspective, SALM-based platforms offer significant potential for advancing biological research, such as studies of stroke pathology, stimulus-induced brain activity, and neurovascular coupling. In summary, SALM opens new possibilities for accessible and scalable neurovascular imaging across platforms and disciplines.

## Methods

### System designs of SALMs

#### Benchtop optical-SALM

The benchtop optical-SALM system utilized two lasers: one operated in CW mode and the other in transistor-transistor logic (TTL) trigger mode. For fluorescent beads imaging (phantom), a 473 nm laser (CW mode, FPYL-473-1000-LED, Frankfurt Laser Company, Germany) and a 488 nm laser (TTL mode, Sapphire LPX, Coherent, USA) were used. For DiD-stained RBC imaging (*in vivo*), we utilized a 660 nm laser (CW mode, gem 660, Laser Quantum, USA) and 640 nm laser (TTL mode, MDL-HD, CNIlaser, China). To ensure homogeneous illumination, a customized dual-fiber bundle (CeramOptec GmbH, Germany) combined the outputs of both lasers into a single beam for epi-illumination of the sample. Fluorescence emission was collected with a camera lens (Laowa Venus 60 mm, Laowa, China), passed through a long-pass filter (FGL515, Thorlabs, USA or LP02-671RU-25, Semrock, USA), and focused on a scientific camera (iXon 888, Andor, U.K.). A multi-channel external trigger (Pulse Pal V2, Sanworks, USA) was employed to synchronize signals, enabling precise timing between pulsed laser illumination and image acquisition. Imaging parameters and trigger signals under three illumination modes and frame rates are shown in Supplementary Note 4.

#### Miniatured optical-SALM

Miniatured optical-SALM was performed using illumination mode 1 and based on the UCLA Miniscope V4 ^[37]^, with an optical configuration consisting of a 470 nm LED excitation source, an excitation filter (ET470/40×, Chroma, USA), a dichroic mirror (T495lpxr, Chroma, USA), an implanted relay GRIN lens (1 mm diameter, 4 mm length, Grintech, Germany), the integrated 0.25-pitch objective GRIN lens assembly, an emission filter (ET525/50 m, Chroma, USA), and a CMOS sensor (MT9V032, On Semiconductor, USA). The relay GRIN lens was chronically implanted in the hippocampus prior to imaging (Supplementary Note 10).

#### NIR-II optical-SALM

NIR-II optical-SALM were performed using illumination Mode 1. An 855 nm CW laser was used to excite the fluorescence signal. Emission was collected using a SWIR camera (WiDy SenS 640V-ST, NiT, France), equipped with a long-pass filter (FELH1100, Thorlabs, USA) and a 50 mm NIR-II camera coated lens (LM50HCSW, Kowa, Japan).

#### Ultrasound-SALM

For transcranial NHP imaging, the transmit frequency was set to 2.23 MHz using a 256-channel phased-array probe (M5ScD, GE Healthcare; nominal center frequency 2.8 MHz) to balance penetration depth, spatial resolution, and MB backscatter amplitude. Multi-angle compounding was performed with 5 full-aperture diverging waves, corresponding to 5 virtual sources uniformly spaced by 5.9 mm and positioned 15.4 mm behind the probe, providing a 70° FOV. Raw data were first corrected for skull-induced aberration and then beamformed using delay-and-sum (DAS) on a polar grid, with radial sampling of λ/2 and an angular step of 0.5°. Tissue clutter and residual noise were subsequently suppressed by singular value decomposition (SVD) filtering on beamformed data to isolate MB signals. For rat imaging after craniotomy, a phased-array of 15.6 MHz center frequency (L22-14vX, Verasonics Inc., Kirkland, WA, USA) was used. Seven plane-wave emissions were compounded for each frame. Image reconstruction procedures were kept the same as described above.

### Phantom experiments

Phantom experiments were performed using an microtubing (inner diameter: 280 µm; outer diameter: 640 µm, #BB31695-PE/1, Scientific Commodities Inc., USA). The tubing was gently bent or knotted to mimic the structural complexity of vascular networks. Orange-yellow fluorescent beads (FMOY-1.3, 460/594 nm, Cospheric, USA) with diameters ranging from 1-5 µm were suspended in water and injected into the tubing. A syringe pump (NE-300, New Era Pump Systems, USA) delivered the fluorescent bead suspension into the tubing at predefined flow rates. Flow velocities were set to 1, 5, 10, and 20 mm/s to mimic physiological blood flow in mouse brain vasculature. For each velocity, image sequences were acquired at 10, 20, and 50 Hz frame rates. Three recordings of 60 s each were collected for each condition. Since the diameter of the microtubing greatly exceeded the resolution limit of the imaging system, super-resolved reconstruction was not performed.

### Animals

#### Cortex imaging with benchtop optical-SALM

A total of N = 9 C57BL/6J mice (12-16 weeks old, female, Charles River Laboratories, Germany) were used for benchtop imaging. Anesthesia was induced via intraperitoneal injection of ketamine (100 mg/kg for induction, 25 mg/kg for maintenance, Pfizer, USA) and xylazine (10 mg/kg for induction, 1.25 mg/kg for maintenance, Bayer, Germany). Buprenorphine (0.1 mg/kg, Bupaq, Streuli Tiergesundheit AG, Switzerland) was administered subcutaneously for analgesia 30 minutes prior to surgery.

For craniotomy, a 3 mm^2^ window was created over the primary somatosensory cortex using a dental drill (Microtorque II, Circuit Medic, USA), and the exposed brain was sealed with a glass coverslip. An oxygen/air mixture (0.2/0.3 L/min) was delivered via a breathing mask to maintain physiological stability, and body temperature was regulated at 37 °C using a feedback-controlled heating system (PhysioSuite, Kent Scientific, USA). Prior to imaging, DiD-stained RBCs (Supplementary Note 7) were injected through the tail vein. A total of 200 s of raw data was recorded, and the super-resolved reconstruction ratio was set to 3. N = 3 mice under craniotomy were assigned for hindpaw stimulation (Fig. 6; Extended Data Fig. 7; Supplementary Note 12), electrical stimuli were applied to the left hind paw with six repetitive stimulation cycles. Each stimulation cycle consisted of 10 s baseline, 8 s stimulation (32 pulses spaced 250 ms), and 22 s post stimulation recording. A trigger generator was used to synchronize the pulsed laser, camera acquisition, and electrical stimulation. A total of 180 s of raw data was recorded with illumination mode 3 and the super-resolved reconstruction ratio was set to 3.

#### Hippocampal imaging with miniaturized optical-SALM

N = 1 C57BL/6J mouse with a chronically implanted GRIN lens in the dorsal hippocampus underwent awake habituation to Miniscope imaging. The Miniscope was secured to a baseplate covering the GRIN lens and returned to its cage for 10 min/day for 3 consecutive days prior to recording. On the day of imaging, the Miniscope was mounted onto the pre-attached baseplate. To minimize animal stress and heating, the 470 nm LED output was limited to ~0.5 mW at the lens tip. The CMOS sensor was set to 752 × 480 pixels at 30 Hz to ensure high SNR and avoid frame loss during continuous acquisition. Fluorescein isothiocyanate (FITC) was first injected to label blood plasma for motion-blur calibration ^[50]^, followed by the injection of fluorescent beads (Fluospheres-F8827, 500/515 nm, Thermo Fisher, USA) to generate streaks. A total of 1000 frames were recorded, and the super-resolved reconstruction ratio was set to 2 due to limited SNR in the miniatured system.

#### Cortex imaging with NIR-II optical-SALM

N = 1 athymic nude-Foxn1nu mouse (5 weeks old, female, Envigo BMS B.V., Netherlands) was used for NIR-II imaging. Anesthesia was induced and maintained with inhaled isoflurane (5% for induction, 1.5% for maintenance) in a gas mixture of 0.1 L/min oxygen and 0.4 L/min air. Data recordings were collected with the scalp intact and after scalp removal, following the same surgical procedures described above. Prior to imaging, microdroplets containing core/shell PbS/CdS quantum dots (NBDY-0038, 1600 nm emission, Nirmidas Biotech, USA; Supplementary Note 11) were administered intravenously. The exposure time was set to 50 ms to ensure sufficient SNR, resulting in an effective frame rate of ~18 Hz. A time-lapse image sequence of 10800 frames was acquired. The super-resolution reconstruction ratio was set to 2 due to limited SNR in the NIR-II system. For comparison, WF images with scalp intact and after scalp removal were collected post intravenous injection of 25 µL aqueous PbS/CdS quantum dots (5 mg/mL in phosphate-buffered saline).

#### Deep brain imaging with ultrasound-SALM

N = 1 adult male rhesus macaque (8 kg, 6 years old) and N = 1 Sprague–Dawley rat with cranial window were secured in a stereotaxic frame to minimize head motion, and body temperature was maintained using a heating blanket. Anesthesia was continuously maintained with 2.5% isoflurane throughout the experiments. For the NHP, the scalp and skull remained intact; only hair was removed to ensure acoustic coupling and reduce coupling-related artifacts. ULM data were acquired in the sagittal (NHP) and coronal (rat) planes. The contrast agent was prepared as a suspension of MBs (SonoVue, Bracco, Milan, Italy) in 5 mL. The total acquisition duration was 180 s for both experiments, and the super-resolution reconstruction ratio was set to 4.

### System validation with high-speed camera data

Optical system validation was performed using data acquired from a high-speed camera (pco.dimax S1, PCO AG, Germany). To simulate reduced frame-rate conditions, datasets originally recorded at 800 Hz were temporally downsampled to 40 Hz by averaging specific subsets of frames, as illustrated in Extended Data Figs. 4-6. All other experimental conditions were consistent with the optical benchtop setup, except for the camera. The total recording time was 120 seconds.

For illumination mode 1, every 20 consecutive frames were averaged to generate one output frame. For mode 2, frames 5, 13, and 14 were removed from each 20-frame block to simulate laser pulse-off intervals, and the remaining frames were averaged. For mode 3, frames 1, 8, and 20 were assigned weighting factors ranging from 2 to 5 to mimic the effect of bright pulsed illumination, after which all frames were averaged to obtain the final image. Structural, velocity and direction maps were reconstructed from the high-frame-rate data using conventional LAT, and served as GT. Quantitative comparisons were performed using PSNR and SSIM. The super-resolved reconstruction ratio was set to 3.

### Localization- and tracking-free neural network

The proposed LTf-Net network is based on the LSTM-UNet architecture ^[30]^, where the bottleneck layers of a conventional UNet are replaced with LSTM units to incorporate temporal context (Supplementary Note 2). This UNet-style design ensures compatibility with varying spatial and temporal input dimensions, making it suitable for data acquired under different illumination modes. Prior to inference, inputs are spatially interpolated according to the desired super-resolution factor, which depends on the SNR. We trained three independent networks to predict super-resolved structural, velocity, and direction maps.

#### Network architecture

The network architecture consists of a UNet-style encoder ^[28]^ for spatiotemporal feature extraction, followed by LSTM units and a symmetric decoder ^[29]^. Each encoder block includes 3×3 convolutions with batch normalization and ReLU, and downsampling via 2×2 max pooling. Feature maps from all input frames are concatenated temporally and processed by convolutional LSTM cells, which aggregate time-series information via learned gating mechanisms. The decoder mirrors the encoder in reverse, using upsampling and convolution layers to reconstruct super-resolved outputs. A final 3×3 convolutional layer produces two output channels for the structural, velocity, or direction map. For direction prediction in the classification setup, the output layer is modified to produce five class logits corresponding to four angular quadrants and one static category.

#### Network training

Training was performed on 10,000 synthetic input–output pairs generated using the developed simulation engine. We used the Adam optimizer with an initial learning rate of 0.003, decayed by 10% every 20 epochs, and trained for 300 epochs. Mean squared error (MSE) loss was used for structural and velocity predictions, while cross-entropy loss was applied for direction classification. Training was conducted on three NVIDIA RTX 4090D GPUs with a batch size of 64, taking approximately 10 hours for each training.

### Simulation engine

The simulation engine (Extended Data Fig. 1) begins by seeding emitters based on biologically inspired vessel density and branching statistics. Each particle is then propagated along tortuous, power-law–distributed trajectories that capture both capillary stalls and bolus flows ^[51]^. At each time step, particles advance by discrete pixel vectors sampled from randomized angles, with slight jitter introduced to avoid unnaturally straight paths. Upon encountering vessel boundaries, particles are redirected stochastically. Each tracer is assigned a Gaussian-distributed brightness value.

To simulate temporal characteristics, the high-resolution streaks are divided into three consecutive temporal bins, each with ± 20% random variation in duration to mimic fine-scale velocity fluctuations within a single exposure. Next, each temporal bin is encoded according to one of three illumination modes. In illumination mode 1, the raw segments are used directly as sequential frames, with inter-frame displacements providing velocity information. In mode 2, two “off” intervals are introduced within each segment to simulate pulsed laser interruptions. In mode 3, bright pulses are superimposed at the beginning, the midpoint, and the end of each segment, dividing the streak into two known temporal intervals for instantaneous flow estimation.

Finally, to replicate real imaging conditions, each bin was spatially downsampled using nearest-neighbor interpolation to emulate detector pixelation, followed by convolution with a Gaussian kernel to simulate the PSF, producing realistic diffraction-limited streaks. Additive Gaussian noise with spatially varying variance was then applied. In parallel, the simulation engine computed per-pixel velocity by integrating particle path lengths over known exposure intervals and generated direction maps based on local displacement vectors, resulting in richly annotated ground truth for both velocity and direction.

### Data analysis for stimulation experiments

Because the imaging FOV contained both fast- and slow-flowing vessels, single-frame reconstructions alone could not reliably capture low-velocity flows, which rarely formed distinct fluorescence streaks. To guide high-temporal-resolution analysis, we first computed flow correlation maps by grouping every three consecutive frames and inputting them into LTf-Net to generate averaged velocity maps at 1 Hz. Pearson correlation coefficients between each pixel’s velocity trace and the mean stimulus-evoked profile identified arteries and veins with strong functional responses (Supplementary Note 12). Vessels with high correlation values were selected for single-frame analysis. Each original frame was then processed individually by LTf-Net to generate a super-resolved velocity map. Non-zero velocity values within the selected vessel regions were averaged per frame and further averaged across six stimulation cycles to produce high-temporal-resolution response curves. For response curve visualization, the mean trace and its standard error of the mean (SEM), calculated from six stimulation cycles at the original high frame rate were presented. The mean trace was additionally smoothed using a Savitzky-Golay filter to enhance the visualization of temporal trends ^[52]^.

